# Cryo-EM confirms a common fibril fold in the heart of four patients with ATTRwt amyloidosis

**DOI:** 10.1101/2024.03.08.582936

**Authors:** Binh An Nguyen, Virender Singh, Shumaila Afrin, Preeti Singh, Maja Pekala, Yasmin Ahmed, Rose Pedretti, Jacob Canepa, Andrew Lemoff, Barbara Kluve-Beckerman, Pawel Wydorski, Farzeen Chhapra, Lorena Saelices

**Affiliations:** Center for Alzheimer’s and Neurodegenerative Diseases, University of Texas Southwestern Medical Center (UTSW), Dallas, TX, USA; Department of Biophysics, University of Texas Southwestern Medical Center (UTSW), Dallas, TX, USA; Peter O’Donnell Jr Brain Institute, University of Texas Southwestern Medical Center (UTSW), Dallas, TX, USA; Department of Biochemistry, University of Texas Southwestern Medical Center, Dallas, TX, USA; Department of Pathology and Laboratory Medicine, Indiana University School of Medicine, Indianapolis, IN, USA

## Abstract

ATTR amyloidosis results from the conversion of transthyretin into amyloid fibrils that deposit in tissues causing organ failure and death. This conversion is facilitated by mutations in ATTRv amyloidosis, or aging in ATTRwt amyloidosis. ATTRv amyloidosis exhibits extreme phenotypic variability, whereas ATTRwt amyloidosis presentation is consistent and predictable. Previously, we found an unprecedented structural variability in cardiac amyloid fibrils from polyneuropathic ATTRv-I84S patients. In contrast, cardiac fibrils from five genotypically-different patients with cardiomyopathy or mixed phenotypes are structurally homogeneous. To understand fibril structure’s impact on phenotype, it is necessary to study the fibrils from multiple patients sharing genotype and phenotype. Here we show the cryo-electron microscopy structures of fibrils extracted from four cardiomyopathic ATTRwt amyloidosis patients. Our study confirms that they share identical conformations with minimal structural variability, consistent with their homogenous clinical presentation. Our study contributes to the understanding of ATTR amyloidosis biopathology and calls for further studies.

**One-Sentence Summary:** Wild-type cardiac ATTR fibrils are structurally homogeneous.

## Introduction

Transthyretin amyloidosis (or ATTR amyloidosis) is a debilitating and progressive disorder caused by the abnormal build-up of transthyretin amyloid (ATTR) that leads to organ failure and death^1^. Transthyretin is primarily produced by the liver and secreted into the systemic circulation as a tetramer^2^. Aging or transthyretin mutations may trigger the dissociation of these tetramers into monomers, and the consequent formation of extracellular amyloid deposition^3^. In wild-type ATTR (ATTRwt) amyloidosis, these amyloids are composed of wild-type transthyretin and the clinical manifestation is primarily cardiomyopathy with late onset^4,5^. In heterozygous variant ATTR (ATTRv) amyloidosis, the deposits contain a mixture of both variant and wild-type transthyretin^6,7^, and the clinical manifestation is extremely variable, presenting a wide spectrum of symptoms such as cardiomyopathy and/or neuropathy, with ocular, gut, and/or kidney involvement, and unpredictable onsets^3,4,8,9^. Homozygous cases of ATTRv amyloidosis often present similar symptomatology as their heterozygous counterparts but with a more rapid disease progression and earlier onsets^10^. The molecular basis of the clinical discrepancies between ATTRwt and ATTRv amyloidosis patients remains elusive.

In amyloidoses of the central nervous system such as tauopathies and synucleinopathies, cryo-electron microscopy (cryo-EM) has revealed an association between the conformation of amyloid fibrils and the disease^11-14^. There are 68 *ex-vivo* structures of human brain amyloid fibrils determined to date (this number will likely be higher by the time of publication of this paper)^12,15^. Most of them feature the wild-type form of the amyloid precursor protein, with three exceptions^12,15^. One exception corresponds to fibrils extracted from the brain of an individual with juvenile-onset synucleinopathy, which contained both wild-type and a mutated form of a-synuclein^16^. The fibril structure in this case differs from those found in wild-type synucleinopathies^14,17^. A second one corresponds to fibrils extracted from two sibling patients that carry the deletion mutation ∆K281 in *MAPT*, leading to Pick’s disease^18^. The fibril structure in this case is identical to that of wild-type tau in Pick’s disease^19^. Worth noting, the tau isoform that is found in these fibrils does not include the R2 repeat, and therefore, the mutated residue^19^. The last exception corresponds to the prion fibril structure isolated from the brain of two *Gerstmann–Sträussler–Scheinker* disease (GSS) patients carrying the F198S mutation^20^. There is, however, no structures determined of wild-type prion fibrils from humans, and thus, their structures cannot be compared. Overall, it is safe to say that all structural studies on wild-type brain fibrils evidence the connection between individual diseases and their specific fibril structures. Inspired by those studies, we hypothesize that the more predictable and consistent phenotypic landscape in ATTRwt amyloidosis is associated with a structural homogeneity of ATTRwt fibrils.

Ten cryo-EM structures of ATTR fibrils have been reported to date^7,21,22^. Cardiac fibrils from five patients with ATTRv-V30M, ATTRv-V20I, ATTRv-G47E, ATTRv-V122I, and ATTRwt genetic backgrounds are structurally homogeneous, sharing the same common core encompassing two fragments^23-25^. In all these structures, the C-terminal fragment folds into a channel made of polar residues that runs throughout the fibril. The remarkable similarity in fibril structures across different ATTR mutations may suggest additional unknown factors influencing the folding events during fibril formation. In contrast, in our recent study we show that cardiac fibrils from three polyneuropathic ATTRv-I84S patients display structural diversity, or polymorphism^22^. We observed that each of the three heterozygous ATTRv-I84S patients present two polymorphs that can co-exist within the same fibril. One polymorph resembles the fold reported in cardiac ATTRwt or ATTRv structures, while the second polymorph shows local variation in the residues that close the polar channel (herein referred to as “the gate”)^22^. We found that these polymorphs with local structural variations are patient-specific. An additional study showed that fibrils extracted from the vitreous humor of an ATTRv-V30M patient are also highly polymorphic, with single, double, and triple filaments. The gate of the polar channel in the double filament ATTRv-V30M fibril structure from the eye adopts a distinct conformation, different from those observed in the other ATTR structures^7,26^. These studies hint towards a possible structural heterogeneity in ATTRv fibrils that depends on phenotype (cardiomyopathy vs neuropathy), the location of fibril deposition, and/or the source of transthyretin since the source of transthyretin in the eye is the retina epithelium instead of the liver. With the exception of ATTRv-I84S structures obtained from three patients, the other ATTR fibril structures were obtained from single patients^22^. As a result, establishing an association between the fibril structure and the clinical presentation in ATTR amyloidosis has been challenging, emphasizing the need to study the fibrils from multiple patients with the same phenotype and genotype.

In this study, we use cryoEM to determine and analyze the structures of *ex-vivo* ATTRwt amyloid fibrils extracted from the hearts of four patients. Our findings confirm the highly structural uniformity in the ordered core of ATTRwt fibrils, sharing the same fold as the previously published ATTRwt fibril structure^24^. They also reveal a fully structured polar channel that appears perturbed in ATTRv-I84S fibrils from the heart and in V30M fibrils from the eye^7,21,22,26^. The structural homogeneity in ATTRwt fibrils may explain the consistent and predictable clinical presentation in patients, thereby suggesting a link between fibril structure and disease presentation, as established from previous observations of brain amyloid fibrils.

## Results

### Extraction and structure determination of ATTRwt fibrils

We successfully extracted the fibrils from the hearts of four patients with ATTRwt amyloidosis (patients ATTRwt-p1, -2, -3 and -4; details in Supplementary Table 1) using a previously published water-based extraction protocol^7,22^. We confirmed the protein identity and genotype using mass spectrometry (Supplementary Fig. 1, Supplementary Table 2). We observed a number of non-tryptic proteoplytic sites in disordered region as well as the amyloid core (Supplementary Fig 2). We also typed the samples using an antibody that recognizes C-terminal fragments of transthyretin and confirmed that the four *ex-vivo* extracts contained type A ATTR fibrils, which are made of both full length and fragments of transthyretin (Supplementary Fig. 3)^27^. For comparison, we used type B ATTR fibrils made of full-length transthyretin and no proteolytic fragments are observed. After extraction and validation, and before data collection, we screened ATTR fibrils for their purity and distribution by negative stain transmission electron microscopy (TEM) and cryo-EM. We found fibrils that appeared to be morphologically uniform in all four cases (Supplementary Fig. 4a). Fibrils were manually picked for helical reconstruction as described in the method section. As observed in our previous study on ATTRv-I84S fibrils, two-dimensional (2D) class averages discerned two fibril polymorphs: straight and curvy^22^. The straight species were unsuitable for structure determination because of their lacking twist. The curvy fibrils with a twisted structure were the most prevalent species (Supplementary Fig. 4b). We stitched together the representative 2D class averages of curvy fibrils and estimated their crossover distances (Supplementary Fig. 4c and Fig. 1). The three-dimensional (3D) class averaging of ATTRwt curvy fibrils resulted in a single class for all four patients with a visually similar structural arrangement of the amyloid core (Supplementary Fig. 4d). The overall resolution of the ATTRwt fibril structures from patients p1, p2, p3, and p4 are 3.31 Å, 3.28 Å, 3.32 Å and 3.43 Å, respectively (Supplementary Table 3).

**Figure 1.**
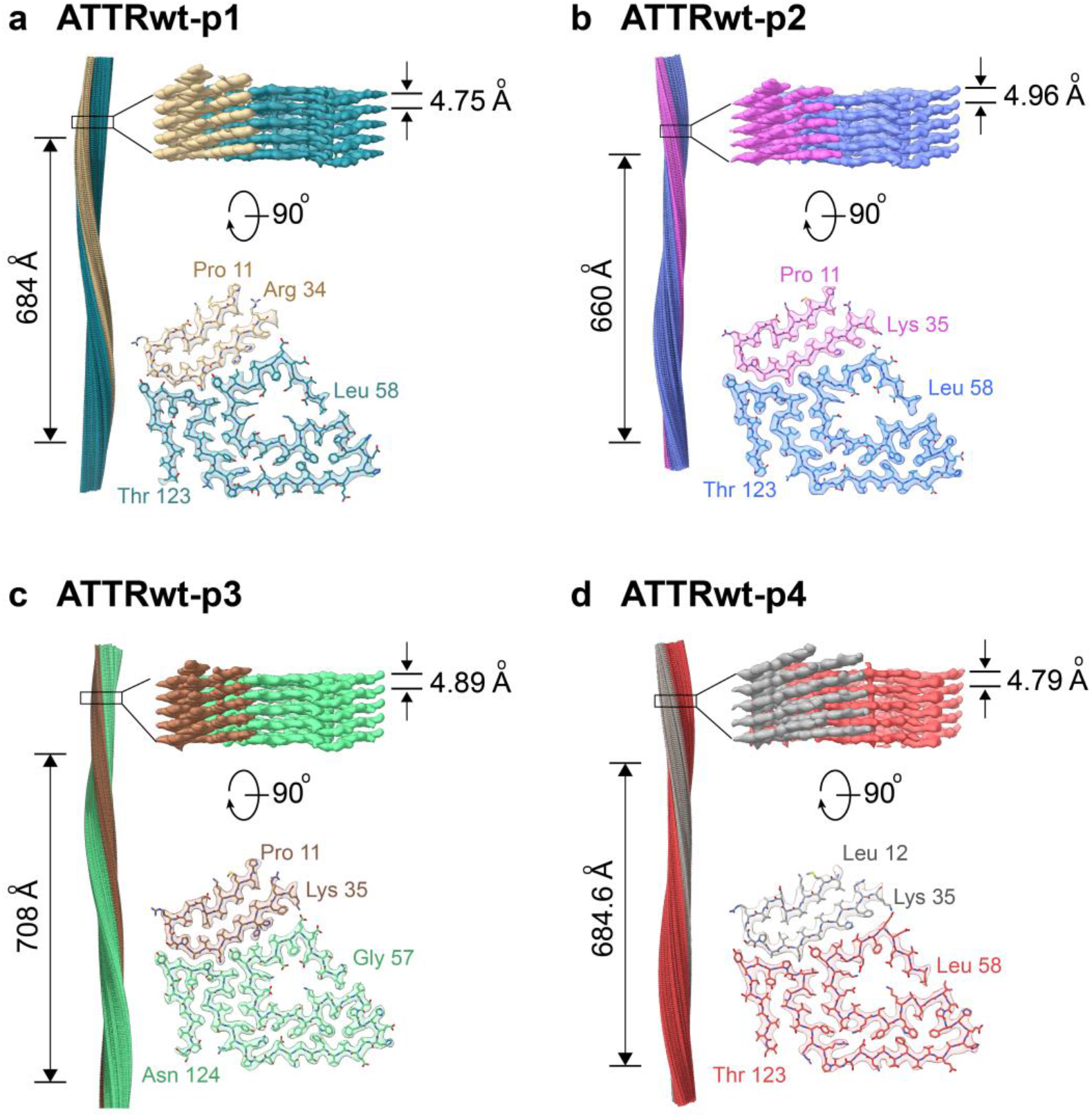
CryoEM density maps and models of ATTRwt fibrils. For each patient (**a** ATTRwt-p1, **b** ATTRwt-p2, **c** ATTRwt-p3 and **d** ATTRwt-p4) figure shows the side view of the reconstructed fibril model with their crossover length (left), the closeup side view of the map with its helical rise (top right), and the top view of a single fibril layer with its model consisting N-terminal fragment (Pro 11/Leu 12 to Arg 34/Lys 35) and C-terminal fragment (Gly 57/Leu 58 to Thr 123/Asn 124) (bottom right.) The color coding is consistent in the cross-sectional model and in the side view of the 3D map.

### Structural commonalities between ATTRwt fibrils

All four ATTRwt fibril structures share similar topology, with one protofilament consisting of two fragments: an N-terminal fragment from Pro 11/Leu 12 to Arg 34/Lys 35 and a C-terminal fragment from Leu 58 to Thr 123 (Fig. 1). The density corresponding to residues Ala 36 to His 56 was missing, suggesting lack or order or their absence. Both ATTRwt-p1 and p3 fibril structures have extra density for His31 rotamers, as previously seen in ATTRwt fibrils^21^, whereas ATTRwt-p2 and p4 fibril structures do not. In contrast to ATTRv-I84S fibrils^22^, the conformation of all ATTRwt fibril structures include a polar channel that resembles a pentagon, starting at Leu 58 and closing at Ile 84. This channel is present in other five ATTR fibril structures from patients with cardiomyopathy or mixed phenotypes^23-25^.

The overall secondary structures of all ATTRwt fibrils including the fibril structure described in the study conducted by Steinebrei et al.^21^ (PDB 8ADE) show similar composition but is different from the native transthyretin (PDB 4TLT) (Fig. 2a). Each fibril layer consists of 14 β-strands (15 β-strands found in three of the four structures (Fig. 2b). The N-terminal fragment consists of three β-strands, with the third β-strand interdigitating with a C-terminal fragment forming what is known as a steric zipper^28^. The C-terminal fragment contains multiple β-strands that run along the fragment, with six of them involved in the formation of the polar channel. A worth noting observation is that only β-strands A, B, F, and H conserve their native secondary structure when converted into fibrils (Fig. 2a). This is of interest because β-strands F and H are the two amyloid-driving segments of transthyretin^29^, and β-strand B appears to be involved in protein oligomerization^30,31^.

**Figure 2.**
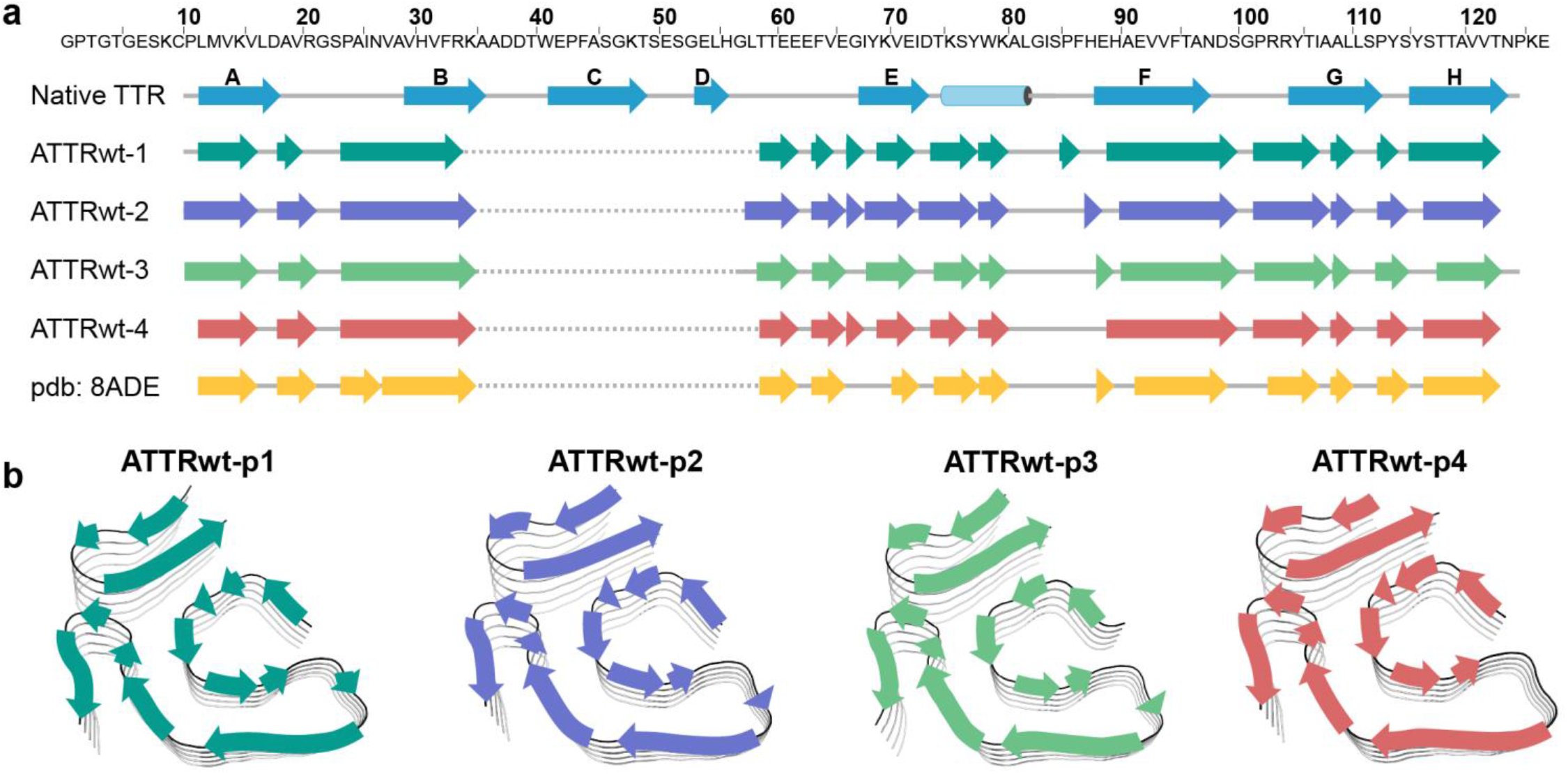
Secondary structure of ATTRwt fibrils. **a** Primary sequence of wild-type transthyretin (top) with secondary structures of native-folded transthyretin (PDB 4TLT) and fibrils from four ATTRwt amyloidosis patients determined in this study (PDB codes 8E7D, 8G9R, 8GBR, and 8E7H) and previous study (PDB 8ADE). **b** Schematic view of the secondary structure on the fibril models of four ATTRwt fibrils.

The solvation energy calculations revealed outstanding estimated stability for ATTRwt fibrils (Fig. 3a), similar to the stability of ATTRv fibrils reported earlier^32^. The estimated solvation energies of the four ATTRwt fibril structures are comparable, both per residue and per chain, averaging -0.68 ± 0.08 kcal/mol and -62.0 ± 6.5 kcal/mol, respectively (Fig. 3a, Supplementary Fig. 5a and Supplementary Table 4). In comparison to the non-ATTR amyloid structures reported in the Amyloid Atlas, and similar to the ATTRv-I84S fibril structures we have recently reported, ATTRwt fibrils are estimated to be more stable, both by residue and chain^15,22,33^. The solvation energy estimates per residue (Fig. 3a and Supplementary Fig. 5a) and residue composition (Fig. 3b and Supplementary Fig. 5b) of ATTRwt fibrils revealed three major hydrophobic pockets contributing to the fibril stability. These pockets encompass the inner interface formed by the hairpin between residues Leu 12 and Val 32 in the N-terminal fragment, the aromatic pocket formed around residues Trp 79 and Phe 95 in the C-terminal fragment, and the triquetra that connects both C- and N-terminal fragments in the center of the structure core (Fig. 3b and Supplementary Fig. 5b). The structure is also stabilized by multiple backbone and side chain interactions, including hydrogen bonding and π-π stacking, as observed in ATTRv fibrils^22^.

**Figure 3.**
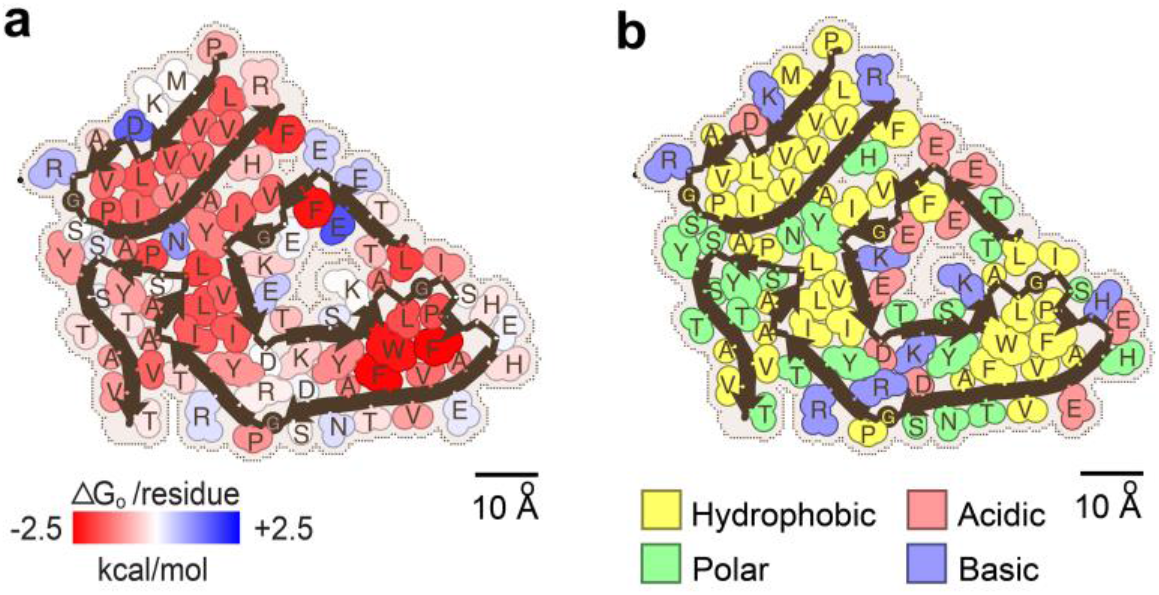
Fibril stability and composition in ATTRwt-p1 amyloidosis. **a** Representation of solvation energies per residue estimated from ATTRwt-p1 fibril structures determined in this study. Residues are colored from favorable (red, -2.5 kcal/mol) to unfavorable stabilization energy (blue, 2.5 kcal/mol). Scale, 10 Å. **b** Schematic view of ATTRwt-p1 fibril structures with residue composition. Residues are color-coded by amino acid category, as labeled. Scale, 10 Å.

### Structural consistency of ATTRwt fibrils

We performed structural alignments of all published ATTR structures (Fig. 4a and 4b) and assessed backbone displacement (only the Cα atoms) through root mean square deviation (RMSD) calculations using GESAMT (Fig. 4c and 4d)^34^. Residues Leu 58 to Glu 66 within the gate region of the polar channel were excluded from analysis due to their contribution to ATTR fibril polymorphism, as previously discussed^22^. The structural alignment confirmed a consistent fold among all ATTRwt fibrils (Fig. 4a). Although the overall alignment of all structures suggests greater structural variability in variants to wild-type, these differences were statistically non-significant (Fig. 4c). When analyzing the RMSD values by residue, ATTRwt-p1, -p3 and -p4 structures exhibit minimal variation with RMSD values through the protein sequence ranging from 0.338 to 0.680 Å. However, the ATTRwt-p2 structure deviates in the N-terminal region and the C-terminal spanning residues Trp79 to Ser100, with an overall RMSD of 1.26 Å (Fig. 4a, c and Supplementary Table 5). The ATTRv structures display greater variability in their backbone structures than ATTRwt structures, spanning from 0.478 to 1.944 (Supplement Table 6), with the highest values for ATTRv-I84S morphologies and the ATTRv-30M fibril structure from the eye (Supplementary Table 6 and Fig. 4b, c, d).

**Figure 4.**
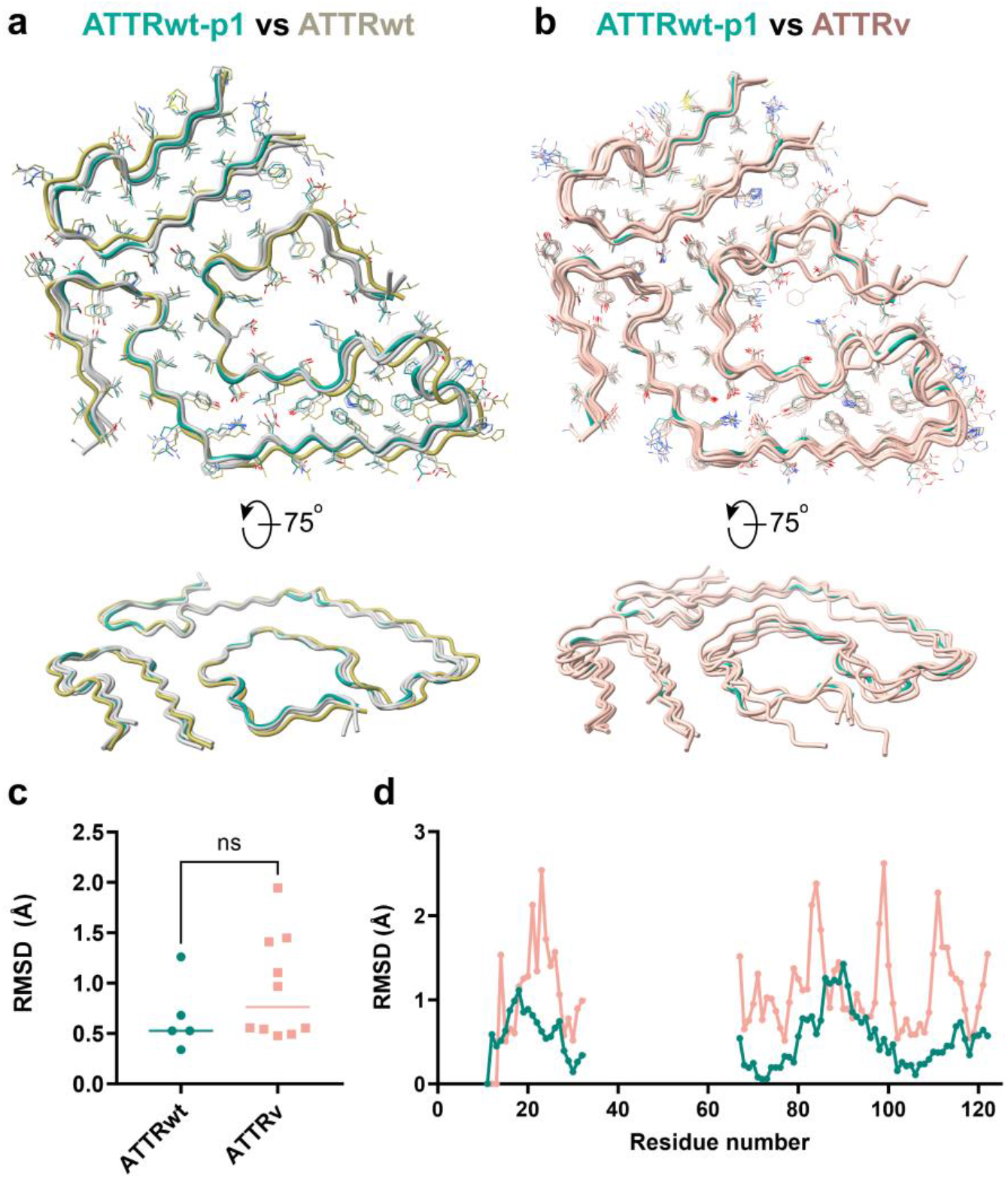
Backbone comparative analysis of ATTR fibril structures. **a** Structure backbone alignment of ATTRwt fibrils with top view (top) and side view (bottom). ATTRwt-p1, in green, is used as a reference, ATTRwt-p2 is in light yellow and three other ATTRwt(s) are in light grey (two from this study and a published ATTRwt PDB 8ADE). **b** Structure backbone alignment of ATTRwt-p1 fibrils, in green; and other published ATTRv structures, in light pink (PDB codes 8E7E, 8E7J, 8E7I, 8TDN, 8TDO, 7OB4, 8PKG, 8PKE, 6SDZ, 8PKF). **c** Overall root mean square deviation (RMSD, only comparing Ca in Å) analysis of the structures included in (**a**) (ATTRwt) and (**b**) (ATTRv only) using GESAMT^34^ Lines indicate mean/median values. **d** Dot plot of the RMSD analysis (only comparing Ca in Å) per residue from the same groups as in (**c**). Each dot represents the average RMSD of comparing to a consensus sequence calculated by GESAMT for each group.

## Discussion

In this study, we used cryo-EM to analyze the structures of *ex-vivo* ATTRwt amyloid fibrils extracted from the hearts of four patients. Our results confirm a highly uniform structure in ATTRwt fibrils, with a consistently ordered amyloid core and symmetry. Despite minor local variations, such as the uncommon His 31 rotamer densities in ATTRwt-p1 and -p2 fibrils, or higher RMSD values of ATTRwt-p2 in comparison to other ATTRwt structures, the overall secondary structure remains identical across all the ATTR-wt fibril structures from various patients and demographics. This indicates a predictable fibril structure in ATTRwt amyloidosis. ATTRwt fibrils also show a range of crossover distances from 660 to 708 Å. Such differences suggest that the amyloid fibrils may accommodate certain level of flexibility without altering the structure core homogeneity, as evidenced by the similarity of secondary structure composition across ATTRwt fibrils.

Our findings add to the question about the potential association between the fibril conformation and disease presentation in ATTR amyloidosis. As observed in the ATTRwt fibrils presented in this manuscript, ATTRv fibrils collected from the heart of multiple patients reveal a common fibril conformation, with the exception of cardiac fibrils extracted from ATTRv-I84S fibrils, which display local structural variations^22^. This raises the question of what differentiate these structurally homogeneous ATTRwt and ATTRv (V30M, V20I, G47E, V122I) fibrils from the heterogenous ATTR-I84S fibrils. There are several potential explanations. One possibility is that the structural variability is mutation-driven, as the mutation from Ile 84 to Ser 84 can disrupt the hydrophobic interaction with Leu58 leading to local variations at the gate of the polar channel^22^. Another explanation is that the structural variability is linked to phenotype. ATTRv-I84S amyloidosis presents predominantly as polyneuropathy, whereas the other genotypes studied so far are associated with cardiomyopathy or a mixed phenotype. If the latter is correct, we anticipate that polyneuropathic genotypes will be associated with structural polymorphism whereas other cardiomyopathic and mixed ATTR genotypes will present a homogeneous structure. The structural homogeneity that we observe in ATTRwt fibrils extracted from four patients is consistent with both explanations. More structural and biological analyses are warranted to get a better understanding of the potential association between phenotype and fibril structure in ATTR amyloidosis.

The little influence of structural polymorphism on the estimated solvation energies and our previous studies on isolated transthyretin peptides inform about a potential aggregation pathway (Fig. 5). Our previous x-ray crystallography study shows that the β-strand B forms out-of-register amyloid fibrils, historically associated with oligomerization^35-39^. This strand and a portion of strand A are part of the N-terminal fragment of the mature ATTR fibril. We speculate that the N-terminal fragment first oligomerizes into a hairpin stabilized by a steric zipper formed in the interior of this hairpin (Fig. 5h). For this to happen, we propose that the segment from residue 36 to 62, approximately, encompassing strand C and D, would initially flip open to expose the N-terminal fragment from residue 11 to 35 (Fig. 5b, c). This proposal is consistent with the previous observation indicating that significant displacements of strands C and D, leads to the disruption of native contacts in the transthyretin structure^40^. This can be facilitated by proteolytic cleavage of residues within this segment, as observed in previous studies and our mass spectrometry analysis (Supplementary Fig. 2), or by mutations that destabilize the native structure^41-44^. In the native structure, the N-terminal fragment adopts a conformation close to a hairpin, so we speculate that the structural reorganization of this segment to convert into the fibril would likely be minimal. The rest of the monomer would then unfurl to open into two wings. The C-terminal fragment from Gly 67 to Thr 123 would reorganize forming the rest of the core (Fig. 5d, e, f, g) followed by the folding of the gate. Notably, there are several cleavage sites that occur at the pivotal points of the fibril structure where Pro and/or Gly are present, suggesting a potential role during protein folding into the final structure of ATTR fibril (Supplementary Fig. 2). We speculate that transthyretin mutations are more likely to affect the folding of the fibrils in the last steps, leading to structural polymorphism. This structural polymorphism could potentially contribute to organ tropism, proteolysis susceptibility, seeding ability, and other features of pathological importance.

**Figure 5.**
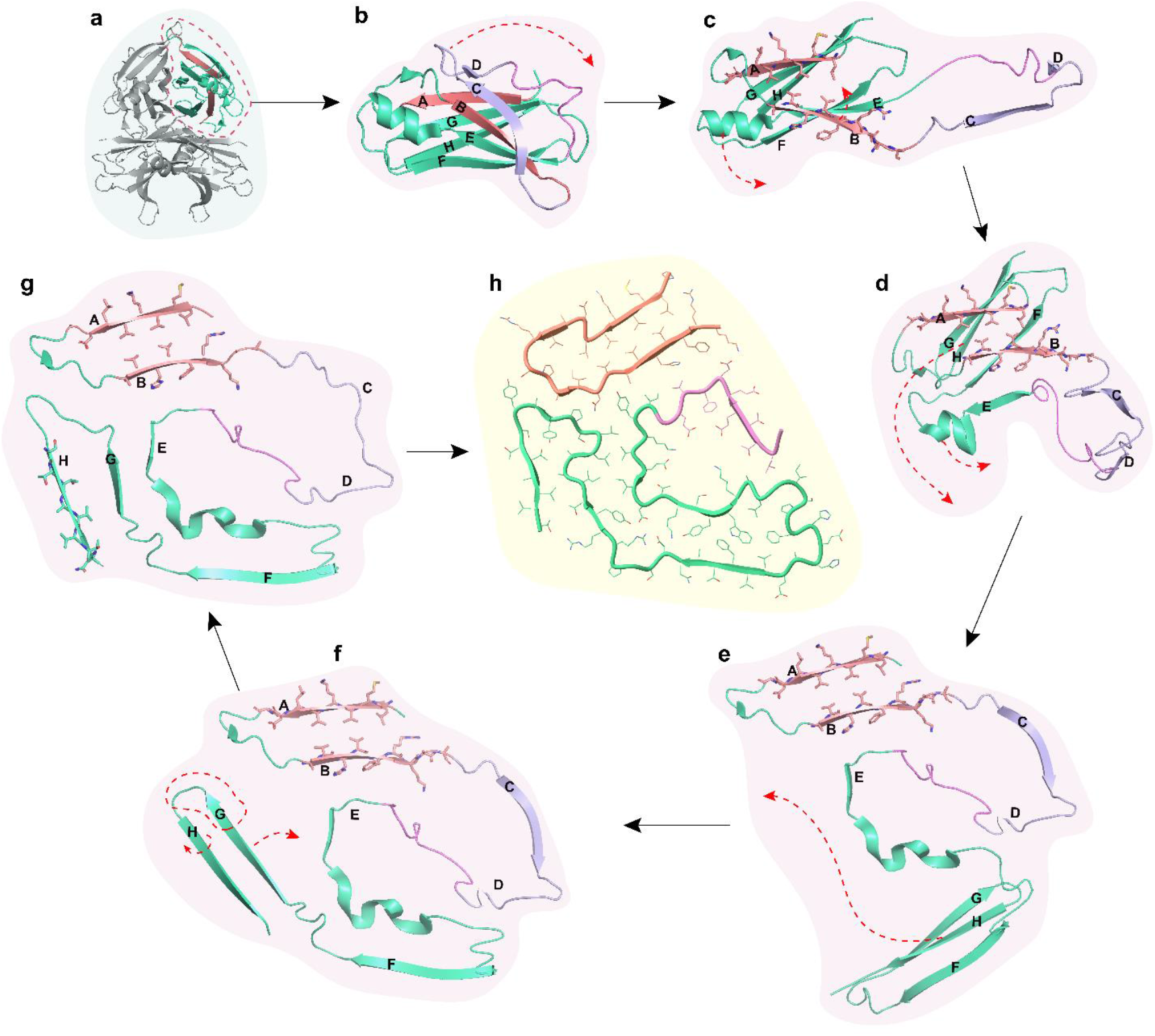
Potential aggregation pathway of transthyretin into ATTR fibrils with minimal structural alteration. **a**, Tetrameric transthyretin (PDB 4TLT). **b**, Monomer unit after dissociating from tetrameric form. **c**, Strands C and D flip away to expose strand A and B. **d**, Strands A and B form hairpin structure while strand E and helix unwind to shape into the core of the polar channel. **e**, Strand G and H rotate to approximately 135° to strand F, so stand F is now free to interact with the back of the polar channel though the formation of an aromatic pocket. **f**, The loop region of strand H and the C-terminal part of strand G unfolds to form the triquetra that connects to the N-terminal hairpin and the back of the polar channel. **g**, Strand H rotates to break the β-sheet hydrogen bonds with strand G and create new interaction at the back of strand G. **h**, Top view of a single layer of the mature ATTR fibril. Color codes represent different regions of the ATTR fibril and monomeric transthyretin. Red, N-terminal hairpin. Purple, unstructured region encompassing strands C and D, which connects the N-terminal to the C-terminal fragments of the ATTR fibril. Pink, the loop between strand D and E that forms the gate of the polar channel. Green, strands E, F, G, and H that form the rest of the C-terminal fragment of the ATTR fibril.

While this study provides important insights into the structure of ATTRwt amyloid fibrils, there is still much to be learned about the structural basis of ATTR amyloidosis. Future research could explore the relationship between fibril structure and disease severity, as well as investigate the mechanisms underlying the early steps in fibril formation and propagation. Together, the understanding of the structural basis of ATTR amyloidosis will enable a more accurate identification of the disease and the identification of novel targets for therapeutics and diagnostics development.

## Methods

### Patients and tissue material

We obtained fresh frozen cardiac tissues from ATTRwt amyloidosis patients (*n* = 4). All samples were postmortem. The details of each tissue sample included in the study are listed in Supplementary Table 1. Specimens from the left ventricle of either explanted or autopsied hearts were obtained from the laboratory of the late Dr. Merrill D. Benson at the University of Indiana. The Office of the Human Research Protection Program granted expedited approval from Internal Review Board review because all specimens were anonymized.

### Extraction of amyloid fibrils from human cardiac tissue

*Ex-vivo* preparations of amyloid fibrils were obtained from the fresh-frozen human tissue as described earlier^7,22^. Briefly, ∼200 mg of frozen cardiac tissue per patient was thawed at room temperature and cut into small pieces with a scalpel. The minced tissue was resuspended into 1 mL Tris-calcium buffer (20 mM Tris, 138 mM NaCl, 2 mM CaCl2, 0.1% NaN3, pH 8.0) and centrifuged for 5 min at 3100 × g and 4 °C. The pellet was washed in Tris-calcium buffer four additional times. After the washing, the pellet was resuspended in 1 mL of 5 mg mL^-1^ collagenase solution (collagenase stock prepared in Tris-calcium buffer) and incubated overnight at 37 °C, shaking at 400 rpm. The resuspension was centrifuged for 30 min at 3100 × g and 4 °C and the pellet was resuspended in 1 mL Tris–ethylenediaminetetraacetic acid (EDTA) buffer (20 mM Tris, 140 mM NaCl, 10 mM EDTA, 0.1% NaN3, pH 8.0). The suspension was centrifuged for 5 min at 3100 × g and 4 °C, and the washing step with Tris–EDTA was repeated nine additional times. All the supernatants were collected for further analysis, when needed. After the washing, the pellet was resuspended in 200 μL ice-cold water supplemented with 5-10 mM EDTA and centrifuged for 5 min at 3100 × g and 4 °C. This step released the amyloid fibrils from the pellet, which were collected in the supernatant. EDTA helped solubilize the fibrils. This extraction step was repeated three additional times. The material from the various patients was handled and analyzed separately.

### Negative-stained transmission electron microscopy

Amyloid fibril extraction was confirmed by transmission electron microscopy as described^22^. Briefly, a 3 μL sample was spotted onto a freshly glow-discharged carbon film 300 mesh copper grid (Electron Microscopy Sciences), incubated for 2 min, and gently blotted onto a filter paper to remove the solution. The grid was negatively stained with 5 µL of 2% uranyl acetate for 2 min and gently blotted to remove the solution. Another 5 μL uranyl acetate was applied onto the grid and immediately removed. An FEI Tecnai 12 electron microscope at an accelerating voltage of 120 kV was used to examine the specimens.

### Western blotting of extracted ATTR fibrils

Western blotting was performed on the extracted fibrils to confirm fibril type. Briefly, 0.5 µg fibrils were dissolved in a tricine SDS sample buffer, boiled for 2 minutes at 85 °C, and run on a Novex™ 16% tris-tricine gel system using a Tricine SDS running buffer. TTR type was determined by transferring the gel contents to a 0.2 µm nitrocellulose membrane and probing with a primary antibody (1:1000) directed against the C-terminal region of the wild type TTR sequence (GenScript). Horseradish peroxidase-conjugated goat anti-rabbit IgG (Invitrogen, at 1:1000) was used as a secondary antibody. Promega Chemiluminescent Substrate (Promega) was used according to the manufacturer’s instructions to visualize TTR content.

### Mass Spectrometry (MS) sample preparation, data acquisition and analysis

For tryptic MS analysis, 0.5 µg of extracted ATTR fibrils were dissolved in a tricine SDS sample buffer, boiled for 2 minutes at 85 °C, and run on a Novex™ 16% tris-tricine gel system using a Tricine SDS running buffer. Gel was stained with Coomassie dye, destained and ATTR smear was cut from the gel. Sample was sent for MS analysis. Samples were digested overnight with trypsin (Pierce) following reduction and alkylation with DTT and iodoacetamide (Sigma–Aldrich). The samples then underwent solid-phase extraction cleanup with an Oasis HLB plate (Waters) and the resulting samples were injected onto an Q Exactive HF mass spectrometer coupled to an Ultimate 3000 RSLC-Nano liquid chromatography system. Samples were injected onto a 75 um i.d., 15-cm long EasySpray column (Thermo) and eluted with a gradient from 0-28% buffer B over 90 min. Buffer A contained 2% (v/v) ACN and 0.1% formic acid in water, and buffer B contained 80% (v/v) ACN, 10% (v/v) trifluoroethanol, and 0.1% formic acid in water. The mass spectrometer operated in positive ion mode with a source voltage of 2.5 kV and an ion transfer tube temperature of 300 °C. MS scans were acquired at 120,000 resolution in the Orbitrap and up to 20 MS/MS spectra were obtained in the ion trap for each full spectrum acquired using higher-energy collisional dissociation (HCD) for ions with charges 2-8. Dynamic exclusion was set for 20 s after an ion was selected for fragmentation.

Raw MS data files were analyzed using Proteome Discoverer v3.0 SP1 (Thermo), with peptide identification performed using a semitryptic search with Sequest HT against the human reviewed protein database from UniProt. Fragment and precursor tolerances of 10 ppm and 0.02 Da were specified, and three missed cleavages were allowed. Carbamidomethylation of Cys was set as a fixed modification, with oxidation of Met set as a variable modification. The false-discovery rate (FDR) cutoff was 1% for all peptides.

The mass spectrometry proteomics data have been deposited to MassIVE data repository, with accession number MSV000094066 ftp://MSV000094066@massive.ucsd.edu.

### Cryo-EM sample preparation, data collection, and processing

Freshly extracted fibril samples were applied to glow-discharged Quantifoil R 1.2/1.3, 300 mesh, Cu grids, blotted with filter paper to remove excess sample, and plunged frozen into liquid ethane using a Vitrobot Mark IV (FEI). Cryo-EM samples were screened on either the Talos Arctica or Glacios at the Cryo-Electron Microscopy Facility (CEMF) at The University of Texas Southwestern Medical Center (UTSW), and the final datasets were collected on a 300 kV Titan Krios microscope (FEI) at the Stanford-SLAC Cryo-EM Center (S^2^C^2^) (Supplementary Table 7). Pixel size, frame rate, dose rate, final dose, and number of micrographs per sample are detailed in Supplementary Table 7. Automated data collection was performed by SerialEM software package^45^ and Thermo Scientific Smart EPU software. The raw movie frames were gain-corrected, aligned, motion-corrected and dose-weighted using RELION’s own implemented motion correction program^46^. Contrast transfer function (CTF) estimation was performed using CTFFIND 4.1^47^. All steps of helical reconstruction, three-dimensional (3D) refinement, and post-process were carried out using RELION 3.1^48^. All filaments were manually picked using EMAN2 e2helixboxer.py^49^. Particles were extracted using a box size of 1024 and 256 pixels with an inter-box distance of 10% of the box length. 2D classification of 1024-pixel particles was used to estimate the helical parameters. 2D classifications of 256-pixel particles were used to select suitable particles for further processing. Fibril helix is assumed to be left-handed. We used an elongated Gaussian blob as an initial reference for 3D classifications of ATTRwt-p1, ATTRwt-p2 and ATTRwt-p4. For ATTRwt-p3, we used a RELION built-in script, *relion_helix_inimodel2d*, to generate an incomplete initial model and low-pass filtered the model to 50 Å. We performed 3D classifications with the average of ∼30k to 40k particles per class to separate filament types. Particles potentially leading to the best reconstructed map were chosen for 3D auto-refinements. CTF refinements and Bayesian polishing were performed to obtain higher resolution. Final maps were post-processed using the recommended standard procedures in RELION. The final subset of selected particles was used for high-resolution gold-standard refinement as described previously^50^. The final overall resolution estimate was evaluated based on the FSC at 0.143 threshold between two independently refined half-maps (Supplementary Fig. 7)^51^.

### Model building

The refined maps were post-processed in RELION before building their models^52^. Our previously published model of ATTRv-P24S (PDB code 8E7I) was used as the template to build the model of ATTR-wt1, which in turn, was used as a template to build the other ATTRwt models. Residue modification, rigid body fit zone, and real space refine zone were performed to obtain the resulting models using COOT^53^. All the statistics are summarized in Supplementary Table 7.

### Stabilization energy calculation

The stabilization energy per residue was calculated by the sum of the products of the area buried for each atom and the corresponding atomic solvation parameters (Supplementary Fig. 5a)^15,22,54^. The overall energy was calculated by the sum of energies of all residues, and assorted colors were assigned to each residue, instead of each atom, in the solvation energy map.

### Statistical Analysis

RMSD data in Cα position per protein structure and per residue were calculated by GESAMT^34,55^. Statistical analysis of fibril stability and RMSD in Cα atoms were performed with Prism 9 for Mac (GraphPad Software) using an unpaired *t* test. All samples were included in the analysis and all measurements are displayed in the graphs.

### Figure Panels

Figure panels were created with ChimeraX 1.6, Pymol 2.3.5 and Adobe Illustrator 2020.

## Acknowledgments

In memory of the Late Dr. Merrill D. Benson, who contributed greatly to the understanding of amyloid diseases and helped affected families for decades. Special thanks to the patients and families who generously donated tissues and the University of Indiana as the source of material. We thank Dr. Michael Sawaya for all of his answers and invaluable advice. We thank the UTSW Cryo-Electron Microscopy Facility, the UTSW Structural Biology Laboratory, the UTSW Electron Microscopy Core Facility, the national cryo-EM facilities Stanford-SLAC (project CA60) and PNCC (project 51267) for instrumentation, technical support, and/or data collection. We thank the UTSW Proteomics core for technical assistance with the proteomics experiments.

## Funding

American Heart Association, Career Development Award 847236, L.S.

National Institutes of Health, National Heart, Lung, and Blood Institute, New Innovator Award DP2-HL163810, L.S.

Welch Foundation, Research Award I-2121-20220331, L.S.

UTSW Endowment, Distinguished Researcher Award from President’s Research Council and start-up funds, L.S.

Cryo-EM research was partially supported by the following grants:

National Institutes of Health grant U24GM129547, Department of Energy Office of Science User Facility sponsored by the Office of Biological and Environmental Research

Department of Energy, Laboratory Directed Research and Development program at SLAC National Accelerator Laboratory, under contract DE-AC02-76SF00515

NIH Common Fund Transformative High Resolution Cryo-Electron Microscopy program (U24 GM129539)

The Cryo-Electron Microscopy Facility and the Structural Biology Laboratory at UTSW are supported by a grant from the Cancer Prevention & Research Institute of Texas (RP170644).

The Electron Microscopy Core Facility at UTSW is supported by the National Institutes of Health (NIH) (1S10OD021685-01A1 and 1S10OD020103-01).

Part of the computational resources were provided by the BioHPC supercomputing facility located in the Lyda Hill Department of Bioinformatics at UTSW. URL: https://portal.biohpc.swmed.edu.

## Author contributions

Conceptualization: B.N., L.S.

Methodology: B.N., V.S., L.S.

Investigation: B.N., V.S. S.A., P.S., M.P., Y.A., R.P., J.C., A.L., B.K., P.W., F.C., L.S.

Visualization: B.N., V.S., L.S.

Funding acquisition: L.S.

Project administration L.S.

Supervision: B.N., V.S., L.S.

Writing – original draft: L.S.

Writing – review & editing: B.N., V.S., S.A., Y.A., R.P., L.S.

## Data and Materials Availability

Structural data have been deposited into the Worldwide Protein Data Bank (wwPDB) and the Electron Microscopy Data Bank (EMDB) with the following EMD accession codes: 26587 (ATTRwt-p1), 29874 (ATTRwt-p2), 29920 (ATTRwt-p3), 26691 (ATTRwt-p4), and PDB accession codes: 8E7D (ATTRwt-p1), 8G9R (ATTRwt-p2), 8GBR (ATTRwt-p3), 8E7H (ATTRwt-p4). The PDB accession code for the previously reported coordinates of ATTRT60A used for data processing is 8E7G. All data generated or analyzed during this study that support the findings are available within this published article and its supplementary data files. Graphed data is provided in the source data file. Cardiac specimens were obtained from the laboratory of the late Dr. Merrill D. Benson at Indiana University. These specimens are under a material transfer agreement with Indiana University and cannot be distributed freely.

