## Supporting Information for "Cryo-EM confirms a common fibril fold in the heart of four patients with ATTRwt amyloidosis"

### Supplementary Figures:

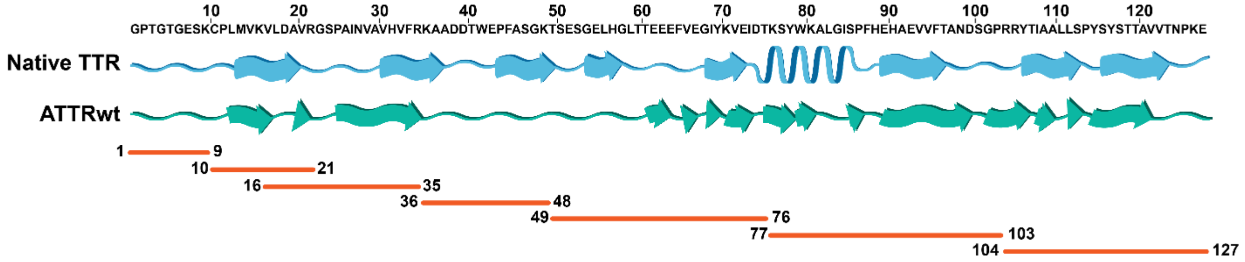


**Supplementary Figure 1:** Results from tryptic mass spectrometry analysis of all ATTRwt samples. We detected all possible tryptic fragments (in orange) covering the whole TTR protein wild-type sequence. The TTR sequence is on the top of the figure; and schematic of the secondary structure of native TTR and ATTRwt fibrils are shown in blue and teal-green respectively.

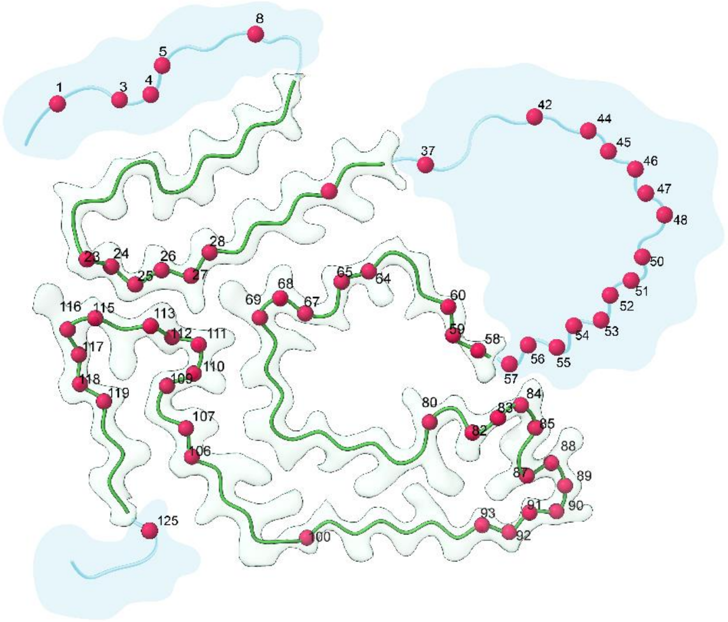


**Supplementary Figure 2.** Schematic of possible proteolytic sites depicted based on non-tryptic peptides from MS data.

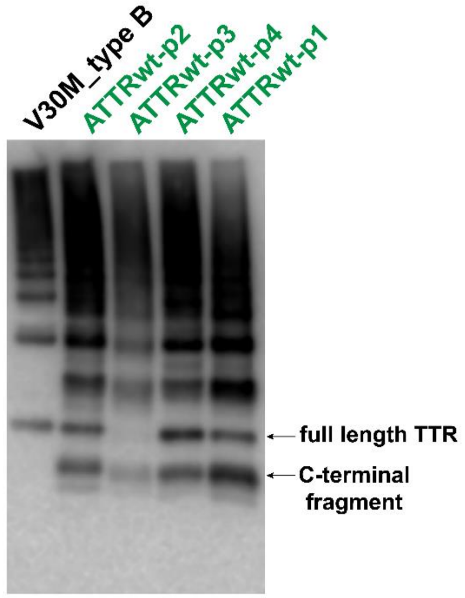


**Supplementary Figure 3:** Western blot analysis of ATTR fibril types from four ATTRwt patients (type A) and a control V30M (type B). The arrows indicate full length TTR (top) and the C-terminal fragment (bottom) which are detected in all fibril types and only in fibril type A, respectively.

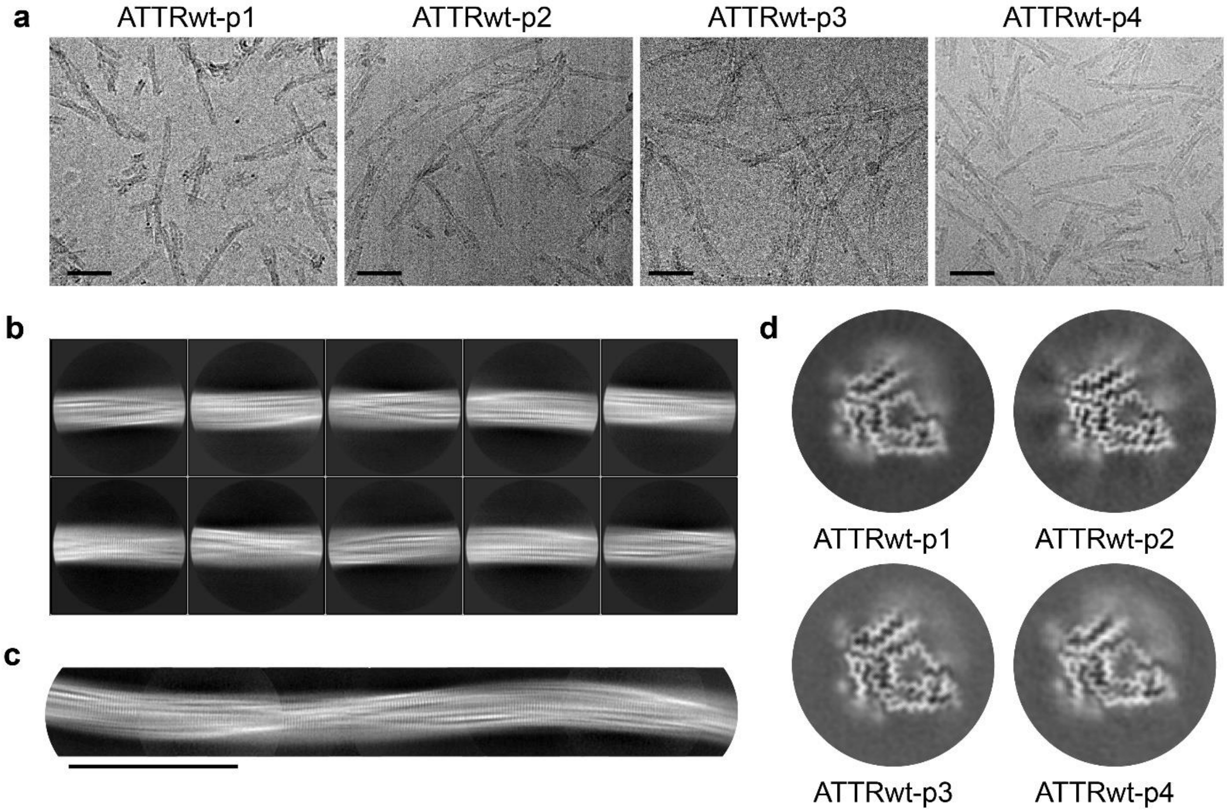


**Supplementary Figure 4. Cryo-EM data collection and processing of cardiac fibrils from four ATTRwt patients (numbered p1 to p4). a** Representative cryo-EM micrographs of ATTRwt fibrils extracted. Scale bars, 100 µm. **b** Representative 2D class averages of curvy fibrils from patient ATTRwt-p1. **c** ATTR curvy fibril after stitching of 2D class averages from patient ATTRwt-p1. Scale bar, 150 Å. **d** 3D class averages of ATTRwt curvy fibrils.

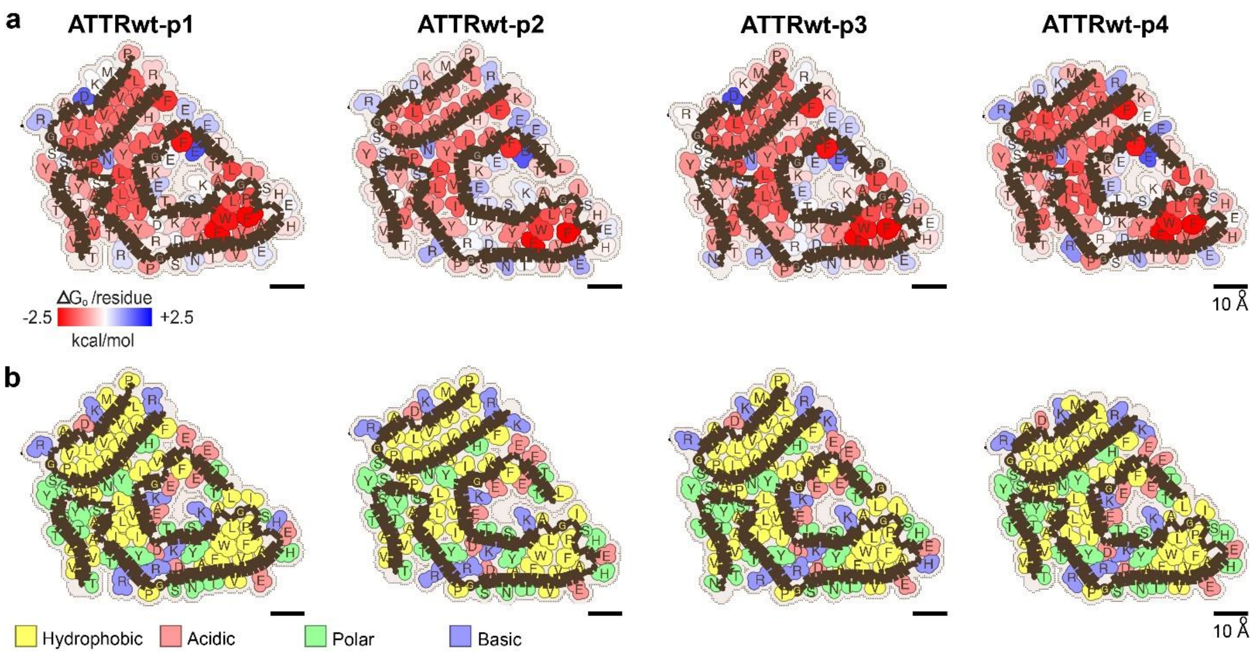


**Supplementary Figure 5. Fibril stability and composition in ATTRwt amyloidosis. a** Representation of solvation energies per residue estimated from ATTRwt fibril structures determined in this study. Residues are colored from favorable (red, -2.5 kcal/mol) to unfavorable stabilization energy (blue, 2.5 kcal/mol). Scale, 10 Å. **b** Schematic view of ATTRwt fibril structures with residue composition. Residues are color-coded by amino acid category, as labeled. Scale, 10 Å. Images for ATTRwt-p1 are also part of Figure 3 in main article.

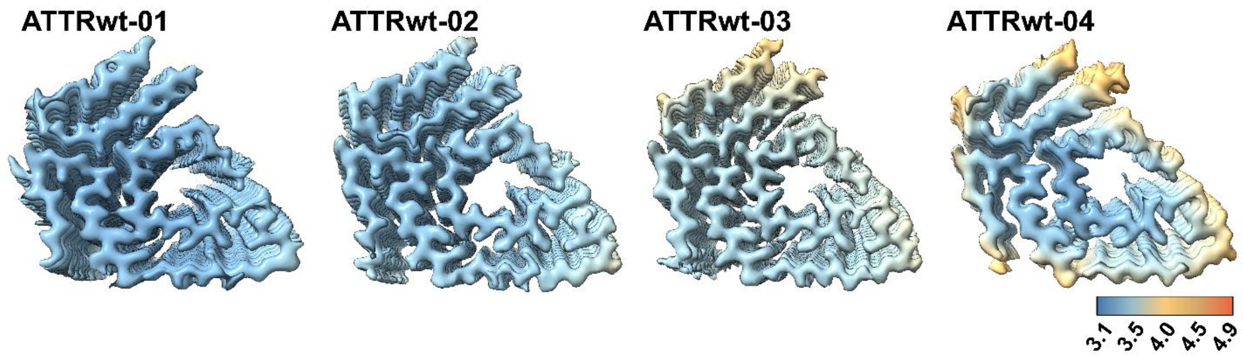


**Supplementary Figure 6. Local resolution maps of four ATTRwt cryo-EM structures.** Local resolution estimation for 3D reconstructions of ATTR fibrils. Resolution scale in Angstroms. Blue represents resolution of 2.8 Å and orange represents 5.6 Å**.**

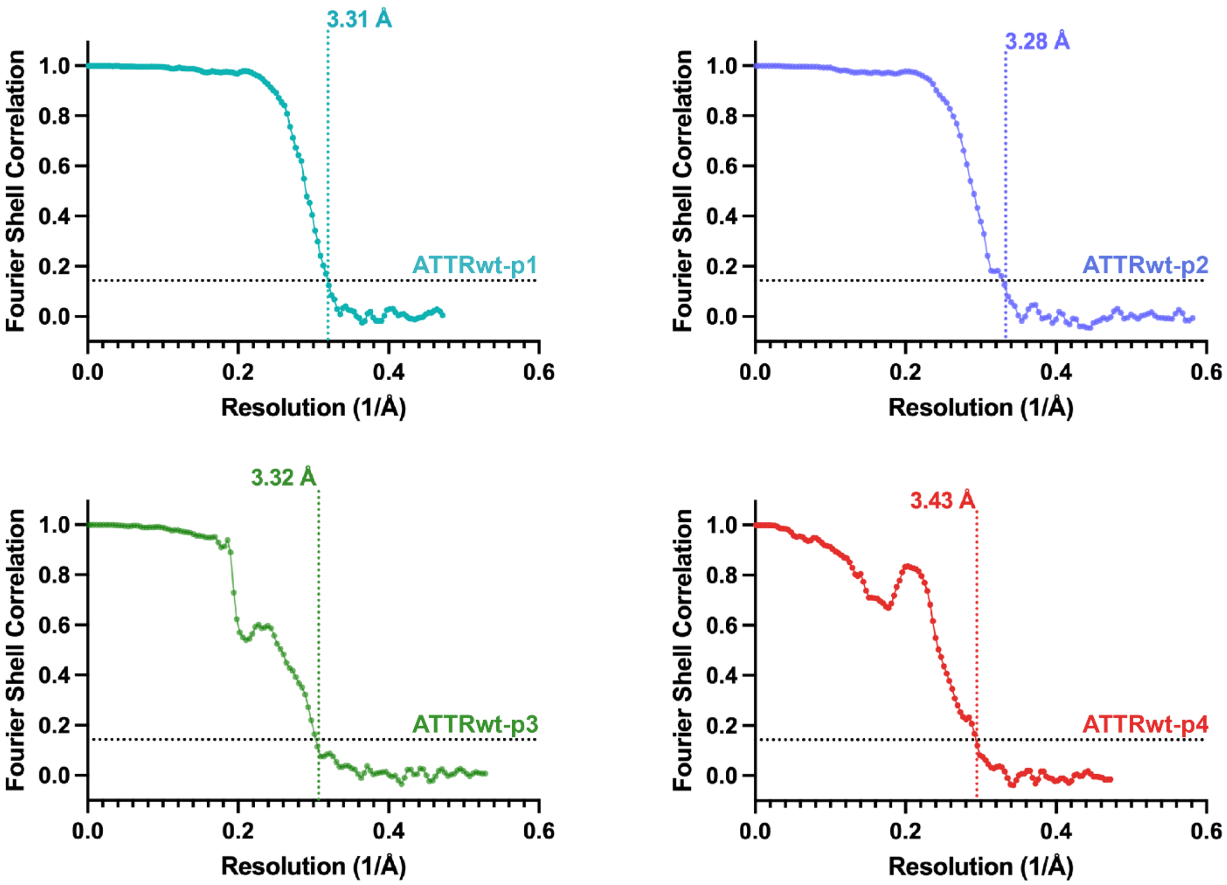


**Supplementary Figure 7. FSC curves.** Evaluation of the resolution of cryo-EM maps by Fourier shell correlation (FSC) curves of two independently refined half-maps from the ATTRwt fibril structures of the four patients.

**Supplementary Tables:**

**Supplementary Table 1.** List of ATTRwt cardiac samples included in the study.

| Genotype | Origin | Sex | Age at collection | Neuropathy signs |
|---|---|---|---|---|
| ATTRwt-p1 | Postmortem | Male | 70 | No, but amyloid found at autopsy |
| ATTRwt-p2 | Postmortem | Male | 83 | No |
| ATTRwt-p3 | Postmortem | Male | 84 | No |
| ATTRwt-p4 | Transplant | Male | 78 | No |

**Supplementary Table 2.** List of Tryptic and Semi-Tryptic Transthyretin peptides [P02766] from MS analysis.

| Annotated Sequence | # PSMs | Positions in Proteins | Abundance ATTRwt-p1 | ATTRwt-p2 | ATTRwt-p3 | ATTRwt-p4 |
|---|---|---|---|---|---|---|
| [A].GPTGTGESK.[C] | 9 | P02766 [21-29] | 3.50E+05 | 1.10E+07 | 8.50E+06 | 2.51E+07 |
| [P].TGTGESKCPLMVK.[V] | 12 | P02766 [23-35] | 1.92E+05 | 3.23E+06 | 6.26E+06 | 5.87E+06 |
| [T].GTGESKCPLMVK.[V] | 9 | P02766 [24-35] | 3.87E+05 | 1.56E+06 | 1.58E+06 | 2.53E+06 |
| [T].GTGESKCPLMVK.[V] | 7 | P02766 [24-35] | 1.37E+05 | 4.72E+05 | 6.12E+05 | 9.96E+05 |
| [G].TGESKCPLMVK.[V] | 10 | P02766 [25-35] | 2.96E+05 | 1.40E+06 | 1.78E+06 | 1.91E+06 |
| [E].SKCPLMVK.[V] | 7 | P02766 [28-35] | 2.34E+05 | 7.56E+05 | 1.86E+06 | 7.31E+05 |
| [K].CPLMVKVLDAVR.[G] | 19 | P02766 [30-41] | 4.84E+05 | 3.09E+06 | 3.55E+06 | 2.48E+06 |
| [K].CPLMVKVLDAVR.[G] | 10 | P02766 [30-41] | 3.11E+05 | 2.03E+06 | 2.03E+06 | 1.63E+06 |
| [K].CPLMVKVLDAVRG.[S] | 10 | P02766 [30-42] | 2.48E+05 | 1.98E+06 | 2.96E+06 | 2.20E+06 |
| [K].VLDAVRGSPAINVAVHVFR.[K] | 31 | P02766 [36-54] | 4.61E+06 | 1.17E+07 | 2.14E+07 | 1.26E+07 |
| [K].VLDAVRGSPAINVAVHVFRK.[A] | 38 | P02766 [36-55] | 5.52E+06 | 1.36E+07 | 1.99E+07 | 1.41E+07 |
| [K].VLDAVRGSPAINVAVHVFRKAADDTWEP.[F] | 33 | P02766 [36-63] | 2.41E+06 | 8.97E+06 | 1.53E+07 | 6.70E+06 |
| [K].VLDAVRGSPAINVAVHVFRKAADDTWEPF.[A] | 34 | P02766 [36-64] | 1.08E+06 | 4.88E+06 | 5.63E+06 | 3.87E+06 |
| [R].GSPAINVAVH.[V] | 14 | P02766 [42-51] | 2.09E+07 | 5.80E+07 | 7.80E+07 | 5.30E+07 |
| [R].GSPAINVAVHV.[F] | 7 | P02766 [42-52] | 2.21E+06 | 1.60E+06 | 1.69E+06 | 7.77E+05 |
| [R].GSPAINVAVHVF.[R] | 19 | P02766 [42-53] | 9.21E+06 | 1.47E+07 | 2.25E+07 | 1.59E+07 |
| [R].GSPAINVAVHVFR.[K] | 758 | P02766 [42-54] | 1.87E+09 | 3.38E+09 | 4.41E+09 | 3.02E+09 |
| [R].GSPAINVAVHVFRK.[A] | 85 | P02766 [42-55] | 3.53E+07 | 1.11E+08 | 1.20E+08 | 7.15E+07 |
| [R].GSPAINVAVHVFRKAADDTWEPFASGK.[T] | 9 | P02766 [42-68] | 1.21E+05 | 1.41E+06 | 5.41E+05 | 2.05E+06 |
| [G].SPAINVAVHVFR.[K] | 53 | P02766 [43-54] | 1.30E+07 | 2.64E+07 | 7.29E+07 | 1.18E+07 |
| [G].SPAINVAVHVFRK.[A] | 7 | P02766 [43-55] | 1.95E+05 | 7.24E+05 | 1.28E+06 | 3.27E+05 |
| [S].PAINVAVHVFR.[K] | 101 | P02766 [44-54] | 1.74E+07 | 3.30E+07 | 3.55E+07 | 2.86E+07 |
| [P].AINVAVHVFR.[K] | 9 | P02766 [45-54] | 1.55E+06 | 3.18E+06 | 2.08E+06 | 2.53E+06 |
| [A].INVAVHVFR.[K] | 18 | P02766 [46-54] | 3.75E+06 | 9.30E+06 | 6.83E+06 | 4.34E+06 |
| [I].NVAVHVFR.[K] | 12 | P02766 [47-54] | 7.75E+05 | 9.83E+05 | 1.55E+06 | 1.14E+06 |
| [R].KAADDTWEP.[F] | 50 | P02766 [55-63] | 5.90E+07 | 1.79E+08 | 4.70E+08 | 1.73E+08 |
| [R].KAADDTWEPF.[A] | 45 | P02766 [55-64] | 4.61E+07 | 1.22E+08 | 2.97E+08 | 9.30E+07 |
| [R].KAADDTWEPFA.[S] | 21 | P02766 [55-65] | 3.87E+06 | 2.60E+07 | 1.79E+07 | 1.77E+07 |
| [R].KAADDTWEPFAS.[G] | 6 | P02766 [55-66] | 2.38E+05 | 2.67E+06 | 2.59E+06 | 2.83E+06 |
| [R].KAADDTWEPFASGK.[T] | 81 | P02766 [55-68] | 4.60E+07 | 3.31E+08 | 3.07E+08 | 4.89E+08 |
| [K].AADDTWEP.[F] | 21 | P02766 [56-63] | 1.07E+07 | 3.33E+07 | 9.57E+07 | 3.28E+07 |
| [K].AADDTWEPF.[A] | 39 | P02766 [56-64] | 7.83E+06 | 1.68E+07 | 3.78E+07 | 1.37E+07 |
| [K].AADDTWEPFA.[S] | 25 | P02766 [56-65] | 1.88E+06 | 1.33E+07 | 9.61E+06 | 1.02E+07 |
| [K].AADDTWEPFASGK.[T] | 60 | P02766 [56-68] | 4.12E+07 | 2.80E+08 | 2.41E+08 | 4.25E+08 |
| [A].ADDTWEPFASGK.[T] | 6 | P02766 [57-68] | 2.51E+04 | 2.61E+05 | 2.67E+05 | 3.76E+05 |
| [D].TWEPFASGK.[T] | 5 | P02766 [60-68] | 1.49E+05 | 5.70E+05 | 7.33E+05 | 1.13E+06 |
| [W].EPFASGK.[T] | 8 | P02766 [62-68] | 1.39E+06 | 1.29E+07 | 4.28E+06 | 6.80E+06 |
| [P].FASGKTSESGELHGLTTEEEFVEGIYK.[V] | 10 | P02766 [64-90] | 1.11E+06 | 1.49E+06 | 4.57E+06 | 1.66E+06 |
| [F].ASGKTSESGELHGLTTEEEFVEGIYK.[V] | 5 | P02766 [65-90] | 5.29E+05 | 2.98E+05 | 6.08E+05 | 3.57E+05 |
| [A].SGKTSESGELHGLTTEEEFVEGIYK.[V] | 16 | P02766 [66-90] | 8.71E+05 | 3.06E+06 | 1.43E+07 | 5.66E+06 |
| [S].GKTSESGELHGLTTEEEFVEGIYK.[V] | 15 | P02766 [67-90] | 2.32E+06 | 5.37E+06 | 2.23E+07 | 1.29E+07 |
| [S].GKTSESGELHGLTTEEEFVEGIYKVEIDTK.[S] | 26 | P02766 [67-96] | 1.37E+06 | 5.63E+06 | 1.47E+07 | 1.16E+07 |
| [G].KTSESGELHGLTTEEEFVEGIYK.[V] | 22 | P02766 [68-90] | 6.88E+06 | 2.40E+07 | 8.77E+07 | 3.35E+07 |
| [G].KTSESGELHGLTTEEEFVEGIYKVEIDTK.[S] | 44 | P02766 [68-96] | 4.89E+06 | 2.50E+07 | 4.70E+07 | 3.03E+07 |
| [K].TSESGELHGLTTE.[E] | 5 | P02766 [69-81] | 3.42E+05 | 8.28E+05 | 1.36E+06 | 8.18E+05 |
| [K].TSESGELHGLTTEEEF.[V] | 8 | P02766 [69-84] | 3.40E+06 | 1.68E+05 | 4.23E+05 | 5.31E+05 |
| [K].TSESGELHGLTTEEEFVEGIYK.[V] | 304 | P02766 [69-90] | 2.80E+08 | 1.04E+09 | 1.85E+09 | 1.39E+09 |
| [K].TSESGELHGLTTEEEFVEGIYKVEID.[T] | 5 | P02766 [69-94] | 1.51E+05 | 2.71E+05 | 2.96E+05 | 6.55E+05 |
| [K].TSESGELHGLTTEEEFVEGIYKVEIDTK.[S] | 577 | P02766 [69-96] | 1.78E+08 | 7.66E+08 | 1.38E+09 | 1.13E+09 |
| [T].SESGELHGLTTEEEFVEGIYK.[V] | 41 | P02766 [70-90] | 9.78E+06 | 2.57E+07 | 1.50E+08 | 5.45E+07 |
| [T].SESGELHGLTTEEEFVEGIYKVEIDTK.[S] | 143 | P02766 [70-96] | 6.50E+06 | 1.68E+07 | 9.39E+07 | 4.04E+07 |
| [S].ESGELHGLTTEEEFVEGIYK.[V] | 24 | P02766 [71-90] | | 2.41E+05 | 3.18E+06 | 8.35E+05 |
| [S].ESGELHGLTTEEEFVEGIYKVEIDTK.[S] | 44 | P02766 [71-96] | 1.28E+06 | 3.37E+06 | 2.10E+07 | 1.09E+07 |
| [E].SGELHGLTTEEEFVEGIYK.[V] | 130 | P02766 [72-90] | 2.31E+07 | 3.36E+07 | 2.06E+08 | 7.31E+07 |
| [E].SGELHGLTTEEEFVEGIYKVEIDTK.[S] | 221 | P02766 [72-96] | 1.90E+07 | 3.32E+07 | 1.81E+08 | 7.86E+07 |
| [S].GELHGLTTEEEFVEGIYK.[V] | 42 | P02766 [73-90] | 1.73E+07 | 1.53E+07 | 6.30E+07 | 2.01E+07 |
| [S].GELHGLTTEEEFVEGIYKVEIDTK.[S] | 102 | P02766 [73-96] | 1.36E+07 | 1.36E+07 | 5.31E+07 | 2.25E+07 |
| [G].ELHGLTTEEEFVEGIYK.[V] | 24 | P02766 [74-90] | 5.02E+05 | 6.64E+05 | 3.41E+06 | 8.14E+05 |
| [G].ELHGLTTEEEFVEGIYKVEIDTK.[S] | 46 | P02766 [74-96] | 5.15E+06 | 6.54E+06 | 2.33E+07 | 1.14E+07 |
| [E].LHGLTTEEEFVEGIYK.[V] | 60 | P02766 [75-90] | 8.80E+06 | 2.18E+07 | 6.02E+07 | 2.44E+07 |
| [E].LHGLTTEEEFVEGIYKVEIDTK.[S] | 99 | P02766 [75-96] | 5.06E+06 | 1.52E+07 | 4.66E+07 | 2.32E+07 |
| [L].HGLTTEEEFVEGIYK.[V] | 17 | P02766 [76-90] | 8.90E+05 | 8.45E+05 | 6.67E+06 | 3.58E+06 |
| [L].HGLTTEEEFVEGIYKVEIDTK.[S] | 14 | P02766 [76-96] | 6.46E+05 | 8.77E+05 | 5.60E+06 | 2.85E+06 |
| [H].GLTTEEEFVEGIYK.[V] | 17 | P02766 [77-90] | 4.75E+06 | 4.16E+06 | 8.91E+06 | 4.19E+06 |
| [H].GLTTEEEFVEGIYKVEIDTK.[S] | 31 | P02766 [77-96] | 4.59E+06 | 5.80E+06 | 1.13E+07 | 6.49E+06 |
| [G].LTTEEEFVEGIYK.[V] | 65 | P02766 [78-90] | 2.55E+07 | 6.01E+07 | 6.83E+07 | 2.51E+07 |
| [G].LTTEEEFVEGIYKVEIDTK.[S] | 210 | P02766 [78-96] | 2.08E+07 | 4.74E+07 | 8.71E+07 | 3.75E+07 |
| [L].TTEEEFVEGIYK.[V] | 9 | P02766 [79-90] | 2.78E+06 | 5.12E+06 | 1.24E+07 | 7.34E+06 |
| [L].TTEEEFVEGIYKVEIDTK.[S] | 19 | P02766 [79-96] | 1.13E+06 | 1.59E+06 | 2.72E+06 | 3.90E+06 |
| [T].TEEEFVEGIYK.[V] | 8 | P02766 [80-90] | 1.13E+06 | 1.92E+06 | 4.99E+06 | 1.65E+06 |
| [T].TEEEFVEGIYKVEIDTK.[S] | 11 | P02766 [80-96] | 2.93E+05 | 4.07E+05 | 1.02E+06 | 5.66E+05 |
| [E].EEFVEGIYK.[V] | 5 | P02766 [82-90] | 4.33E+05 | 4.30E+05 | 1.72E+06 | 8.65E+05 |
| [E].FVEGIYK.[V] | 10 | P02766 [84-90] | 4.95E+06 | 1.71E+07 | 3.36E+07 | 1.55E+07 |
| [F].VEGIYKVEIDTK.[S] | 14 | P02766 [85-96] | 3.60E+06 | 8.80E+04 | 7.93E+05 | 8.02E+05 |

| | | | | | | |
|---|---|---|---|---|---|---|
| [G].IYKVEIDTK.[S] | 9 | P02766 [88-96] | 8.11E+06 | 5.72E+07 | 2.06E+07 | 7.93E+06 |
| [K].VEIDTKSYWK.[A] | 5 | P02766 [91-100] | 2.37E+05 | 7.35E+05 | 1.51E+06 | 7.73E+05 |
| [K].SYWKALGISPFHEHAEVVFTANDSGPR.[R] | 48 | P02766 [97-123] | 2.85E+06 | 1.10E+07 | 1.15E+07 | 8.50E+06 |
| [W].KALGISPFHEHAEVVFTANDSGPR.[R] | 8 | P02766 [100-123] | 1.87E+05 | 7.17E+05 | 5.65E+05 | 7.71E+05 |
| [K].ALGISPFH.[E] | 5 | P02766 [101-108] | 1.44E+06 | 3.18E+06 | 6.46E+06 | 3.23E+06 |
| [K].ALGISPFHE.[H] | 7 | P02766 [101-109] | 1.26E+06 | 2.19E+06 | 5.46E+06 | 3.04E+06 |
| [K].ALGISPFHEHA.[E] | 9 | P02766 [101-111] | 9.17E+05 | 2.31E+06 | 6.70E+06 | 3.02E+06 |
| [K].ALGISPFHEHAEVVFTA.[N] | 5 | P02766 [101-117] | 3.52E+05 | 5.80E+05 | 1.57E+06 | 1.06E+06 |
| [K].ALGISPFHEHAEVVFTAN.[D] | 42 | P02766 [101-118] | 2.22E+06 | 5.77E+06 | 1.21E+07 | 7.72E+06 |
| [K].ALGISPFHEHAEVVFTAND.[S] | 22 | P02766 [101-119] | 1.05E+06 | 1.06E+06 | 3.35E+06 | 2.31E+06 |
| [K].ALGISPFHEHAEVVFTANDS.[G] | 11 | P02766 [101-120] | 2.95E+05 | 5.83E+05 | 1.42E+06 | 9.97E+05 |
| [K].ALGISPFHEHAEVVFTANDSG.[P] | 9 | P02766 [101-121] | 2.19E+05 | 4.17E+05 | 8.07E+05 | 7.27E+05 |
| [K].ALGISPFHEHAEVVFTANDSGPR.[R] | 1253 | P02766 [101-123] | 5.21E+08 | 1.61E+09 | 3.03E+09 | 1.64E+09 |
| [K].ALGISPFHEHAEVVFTANDSGPRR.[Y] | 35 | P02766 [101-124] | 3.42E+06 | 1.61E+07 | 1.43E+07 | 2.85E+07 |
| [A].LGISPFHEHAEVVFTANDSGPR.[R] | 8 | P02766 [102-123] | 4.85E+05 | 1.25E+06 | 2.67E+06 | 1.21E+06 |
| [L].GISPFHEHAEVVFTANDSGPR.[R] | 38 | P02766 [103-123] | 1.68E+06 | 6.37E+06 | 5.17E+06 | 4.89E+06 |
| [G].ISPFHEHAEVVFTANDSGPR.[R] | 17 | P02766 [104-123] | 1.76E+06 | 6.28E+06 | 3.45E+06 | 1.69E+06 |
| [I].SPFHEHAEVVFTANDSGPR.[R] | 5 | P02766 [105-123] | 1.19E+05 | 6.21E+05 | 4.72E+05 | 3.91E+05 |
| [P].FHEHAEVVFTANDSGPR.[R] | 5 | P02766 [107-123] | 9.61E+04 | 5.89E+05 | 2.28E+05 | |
| [F].HEHAEVVFTANDSGPR.[R] | 44 | P02766 [108-123] | 2.65E+06 | 6.26E+06 | 4.82E+06 | 2.27E+06 |
| [H].EHAEVVFTANDSGPR.[R] | 9 | P02766 [109-123] | 5.40E+05 | 8.90E+05 | 1.07E+06 | 7.95E+05 |
| [H].AEVVFTANDSGPR.[R] | 30 | P02766 [111-123] | 1.94E+06 | 2.67E+06 | 3.37E+06 | 2.61E+06 |
| [A].EVVFTANDSGPR.[R] | 7 | P02766 [112-123] | 5.12E+05 | 1.17E+06 | 1.71E+06 | 1.25E+06 |
| [E].VVFTANDSGPR.[R] | 9 | P02766 [113-123] | 1.91E+06 | 1.04E+07 | 5.05E+06 | 2.54E+06 |
| [R].RYTIAALLSPY.[S] | 43 | P02766 [124-134] | 4.09E+07 | 8.17E+07 | 2.02E+08 | 9.88E+07 |
| [R].RYTIAALLSPYS.[Y] | 7 | P02766 [124-135] | 8.61E+05 | 8.68E+05 | 3.57E+06 | 1.74E+06 |
| [R].RYTIAALLSPYSY.[S] | 12 | P02766 [124-136] | 1.37E+07 | 2.32E+07 | 8.58E+07 | 3.92E+07 |
| [R].RYTIAALLSPYSYS.[T] | 8 | P02766 [124-137] | 5.01E+05 | 6.32E+05 | 2.94E+06 | 1.06E+06 |
| [R].RYTIAALLSPYSYST.[T] | 7 | P02766 [124-138] | 4.36E+05 | 3.74E+05 | 1.98E+06 | 3.50E+06 |
| [R].RYTIAALLSPYSYSTT.[A] | 7 | P02766 [124-139] | 6.59E+05 | 4.39E+05 | 3.08E+06 | 2.18E+06 |
| [R].RYTIAALLSPYSYSTTA.[V] | 5 | P02766 [124-140] | 3.90E+05 | 4.77E+05 | 2.92E+06 | 1.59E+06 |
| [R].RYTIAALLSPYSYSTTAVV.[T] | 5 | P02766 [124-142] | 2.85E+05 | 1.65E+05 | 4.23E+05 | 2.26E+05 |
| [R].RYTIAALLSPYSYSTTAVVTN.[P] | 444 | P02766 [124-144] | 1.38E+08 | 1.22E+08 | 1.71E+08 | 1.45E+08 |
| [R].RYTIAALLSPYSYSTTAVVTNPK.[E] | 191 | P02766 [124-146] | 1.20E+08 | 4.39E+08 | 6.57E+08 | 4.29E+08 |
| [R].RYTIAALLSPYSYSTTAVVTNPKE.[-] | 488 | P02766 [124-147] | 5.73E+08 | 1.79E+09 | 2.76E+09 | 1.87E+09 |
| [R].YTIAALLSPY.[S] | 16 | P02766 [125-134] | 7.79E+06 | 1.34E+07 | 3.55E+07 | 1.71E+07 |
| [R].YTIAALLSPYSY.[S] | 13 | P02766 [125-136] | 3.08E+06 | 4.44E+06 | 1.84E+07 | 7.66E+06 |
| [R].YTIAALLSPYSYSTTAVVTN.[P] | 118 | P02766 [125-144] | 7.12E+07 | 5.85E+07 | 1.94E+08 | 9.08E+07 |
| [R].YTIAALLSPYSYSTTAVVTNPK.[E] | 82 | P02766 [125-146] | 5.88E+07 | 2.03E+08 | 4.25E+08 | 2.37E+08 |
| [R].YTIAALLSPYSYSTTAVVTNPKE.[-] | 383 | P02766 [125-147] | 3.12E+08 | 1.02E+09 | 2.11E+09 | 1.50E+09 |
| [Y].TIAALLSPYSYSTTAVVTNPK.[E] | 12 | P02766 [126-146] | 8.76E+05 | 1.67E+06 | 4.10E+06 | 1.64E+06 |
| [Y].TIAALLSPYSYSTTAVVTNPKE.[-] | 21 | P02766 [126-147] | 2.73E+06 | 7.40E+06 | 3.16E+07 | 9.57E+06 |
| [T].IAALLSPYSYSTTAVVTNPKE.[-] | 9 | P02766 [127-147] | | 3.24E+05 | 1.47E+06 | 4.58E+05 |
| [A].ALLSPYSYSTTAVVTNPK.[E] | 6 | P02766 [129-146] | 3.85E+05 | 9.64E+05 | 1.43E+06 | 8.10E+05 |
| [A].ALLSPYSYSTTAVVTNPKE.[-] | 16 | P02766 [129-147] | 2.66E+06 | 6.40E+06 | 1.67E+07 | 7.00E+06 |
| [A].LLSPYSYSTTAVVTNPKE.[-] | 8 | P02766 [130-147] | 4.23E+05 | 1.35E+06 | 3.46E+06 | 1.71E+06 |
| [L].LSPYSYSTTAVVTNPK.[E] | 10 | P02766 [131-146] | 4.63E+05 | 3.04E+06 | 2.36E+06 | 1.35E+06 |
| [L].LSPYSYSTTAVVTNPKE.[-] | 14 | P02766 [131-147] | 2.66E+06 | 1.44E+07 | 1.98E+07 | 1.11E+07 |
| [L].SPYSYSTTAVVTNPKE.[-] | 12 | P02766 [132-147] | 2.48E+05 | 1.69E+06 | 2.91E+06 | 1.29E+06 |
| [S].PYSYSTTAVVTNPKE.[-] | 7 | P02766 [133-147] | 1.01E+05 | 7.01E+05 | 1.74E+06 | 1.00E+06 |
| [Y].SYSTTAVVTNPK.[E] | 15 | P02766 [135-146] | 8.96E+06 | 4.40E+07 | 5.18E+07 | 2.68E+07 |
| [Y].SYSTTAVVTNPKE.[-] | 34 | P02766 [135-147] | 3.74E+07 | 2.05E+08 | 3.57E+08 | 1.50E+08 |
| [S].YSTTAVVTNPKE.[-] | 5 | P02766 [136-147] | 1.68E+05 | 7.69E+05 | 1.60E+06 | 8.09E+05 |
| [Y].STTAVVTNPK.[E] | 8 | P02766 [137-146] | 3.00E+06 | 1.11E+07 | 1.50E+07 | 8.46E+06 |
| [Y].STTAVVTNPKE.[-] | 17 | P02766 [137-147] | 1.70E+07 | 7.15E+07 | 1.35E+08 | 7.60E+07 |
| [S].TTAVVTNPKE.[-] | 6 | P02766 [138-147] | 3.09E+05 | 9.32E+05 | 2.55E+06 | 1.52E+06 |
| [T].TAVVTNPKE.[-] | 8 | P02766 [139-147] | 4.05E+05 | 7.04E+05 | 1.87E+06 | 4.82E+06 |

**Supplementary Table 3.** Summary of overall, maximum, and minimum resolution values of ATTR fibril maps shown in Figure

| **Resolution (Å)** | **ATTRwt-p1** | **ATTRwt-p2** | **ATTRwt-p3** | **ATTRwt-p4** |
|---|---|---|---|---|
| Overall | 3.31 | 3.28 | 3.32 | 3.43 |
| Maximum | 3.08 | 3.18 | 3.1 | 3.3 |
| Minimum | 3.37 | 3.8 | 4.2 | 4.6 |

**Supplementary Table 4.** Stabilization energies per residue and per chain of ATTRwt fibrils in kcal/mol.

| | **ATTRwt-p1** | **ATTRwt-p2** | **ATTRwt-p3** | **ATTRwt-p4** |
|---|---|---|---|---|
| Per chain | -67.3 | -51.3 | -66.0 | -65.5 |
| Per residue | -0.75 | -0.56 | -0.71 | -0.73 |

**Supplementary Table 5**. Root mean square deviation (RMSD of Cα atoms) of ATTRwt structures against a consensus RMSD calculated by GESAMT.

| **Structures** | **RMSD** (Å) |
|---|---|
| ATTRwt-p1 | 0.527 |
| ATTRwt-p2 | 1.26 |
| ATTRwt-p3 | 0.338 |
| ATTRwt-p4 | 0.527 |
| ATTRwt, PDB 8ADE | 0.68 |

**Supplementary Table 6**. Root mean square deviation (RMSD of Cα atoms) of ATTRv structures against a consensus RMSD calculated by GESAMT.

| **Structures** | | **RMSD (Å)** |
|---|---|---|
| ATTRv-I84S, Absent gate, | PDB 8E7E | 1.944 |
| ATTRv-I84S, Broken gate, | PDB 8E7J | 1.409 |
| ATTRv-I84S, Open gate, | PDB 8TDO | 0.966 |
| ATTRv-I84S, Closed gate, | PDB 8TDN | 0.491 |
| ATTRv-P24S, | PDB 8E7I | 1.448 |
| ATTRv-V20I, | PDB 8PKE | 0.543 |
| ATTRv-V122I, | PDB 8PKG | 0.556 |
| ATTRv-G47E, | PDB 8PKF | 0.478 |
| ATTRv-V30M, eye, | PDB 7OB4 | 1.103 |
| ATTRv-V30M, | PDB 6SDZ | 0.553 |

**Supplementary Table 7.** Data collection and refinement statistics.

| Data collection | ATTRwt-p1 | ATTRwt-p2 | ATTRwt-p3 | ATTRwt-p4 |
|---|---|---|---|---|
| **Microscope** | Titan Krios | Titan Krios | Titan Krios (G3i) | Titan Krios (G3i) |
| **Acceleration Voltage (kV)** | 300 | 300 | 300 | 300 |
| **Detector** | K3 | K3 | K3 | K3 |
| **Software** | SerialEM 3.8 | EPU | EPU | SerialEM 3.8 |
| **Magnification** | 165,000x | 105,000x | 105,000 | 165,000x |
| **Pixel size at detector (Å/px)** | 0.53 | 0.86 | 0.946 | 0.53 |
| **Defocus range (μm)** | 1.0 to 2.2 | 0.9 to 2.5 | 1.2 to 2.2 | 1.0 to 2.2 |
| **Total dose (e)** | 46 | 50 | 40 | 46 |
| **Exposure time (sec)** | 2.4 | 1.81 | 4.98 | 2.4 |
| **Number of movie frames** | 51 | 40 | 40 | 51 |
| **Usable micrograph** | 3295 | 5051 | 3104 | 2698 |
| **Box size (pixel)** | 256 | 256 | 256 | 256 |
| **Total extracted segments** | 316273 | 252357 | 275727 | 388803 |
| **Number of segments after 2D (curvy)** | 151781 | 192267 | 151524 | 278654 |
| **Number of straight segments** | 20657 | 15833 | 19121 | 58116 |
| **Number of segments after 3D** | 80229 | 37947 | 50584 | 11420 |
| **Symmetry imposed** | C1 | C1 | C1 | C1 |
| **Helical rise (Å)** | 4.75 | 4.96 | 4.90 | 4.79 |
| **Helical twist (°)** | 1.25 | 1.35 | 1.245 | 1.26 |
| **Crossover length (Å)** | 684 | 661.3 | 708.4 | 684 |
| **B factor** | -79.23 | -93.5 | -132.8 | -54.8 |
| **Map resolution (Å; FSC=0.143)** | 3.31 | 3.28 | 3.32 | 3.43 |
| **Map resolution (Å; FSC=0.5)** | 3.68 | 3.64 | 3.77 | 4.17 |
| **Total number of straight filaments** | 3716 | 33487 | 13416 | 82211 |
| **Total number of curvy filaments** | 126640 | 192267 | 151524 | 278654 |
| **Straight fibril particles (%)** | 2.93 | 14.8 | 8.13 | 22.3 |
| **Non-hydrogen atoms** | 3525 | 3570 | 3630 | 3535 |
| **Protein residues** | 450 | 455 | 465 | 450 |
| **Number of chains** | 5 | 5 | 5 | 5 |
| **Water/ligands** | 0/0 | 0/0 | 0/0 | 0/0 |
| **MolProbity score** | 2.33 | 1.42 | 1.11 | 2.25 |
| **Clash score** | 21.89 | 2.94 | 3.17 | 23.62 |
| **Rotamer outliers (%)** | 0.26 | 0 | 0 | 0 |
| **R.M.S deviations bonds (Å)** | 0.005 | 0.002 | 0.004 | 0.004 |
| **R.M.S deviations angle (°)** | 0.703 | 0.503 | 0.593 | 0.648 |
| **Ramachandran plot** | | | | |
| Favored | 91.86 | 95.17 | 98.88 | 94.42 |
| Allowed | 8.14 | 4.83 | 1.12 | 5.58 |
| Outliers | 0 | 0 | 0 | 0 |
| **CaBLAM outliers (%)** | 2.44 | 3.61 | 5.88 | 2.44 |
| **Model vs Data** | 0.76 | 0.86 | 0.86 | 0.78 |
